# Multiplexed confocal and super-resolution fluorescence imaging of cytoskeletal and neuronal synapse proteins

**DOI:** 10.1101/111625

**Authors:** Syuan-Ming Guo, Remi Veneziano, Simon Gordonov, Li Li, Demian Park, Anthony B. Kulesa, Paul C. Blainey, Jeffrey R. Cottrell, Edward S. Boyden, Mark Bathe

**Affiliations:** Department of Biological Engineering, MIT, Cambridge, MA, USA; Stanley Center for Psychiatric Research, Broad Institute of MIT and Harvard, Cambridge, MA, USA; Media Lab, MIT, Cambridge, MA, USA; Broad Institute of MIT and Harvard, Cambridge, MA, USA; McGovern Institute for Brain Research, Department of Brain and Cognitive Sciences, MIT, Cambridge, MA, USA

## Abstract

Neuronal synapses contain dozens of protein species whose expression levels and localizations are key determinants of synaptic transmission and plasticity. The spectral properties of fluorophores used in conventional microscopy limit the number of measured proteins to four species within a given sample. The ability to perform high-throughput confocal or super-resolution imaging of many proteins simultaneously without limitation in target number imposed by this spectral limit would enable large-scale characterization of synaptic protein networks in situ. Here, we introduce PRISM: Probe-based Imaging for Sequential Multiplexing, a method that sequentially utilizes either high affinity Locked Nucleic Acid (LNA) or low affinity DNA probes to enable diffraction-limited confocal and PAINT-based super-resolution imaging. High-affinity LNA probes offer high-throughput, confocal-based imaging compared with PAINT, which uses low affinity probes to realize localization-based super-resolution imaging. Simultaneous immunostaining of all targets is performed prior to imaging, followed by sequential LNA/DNA probe exchange that requires only minutes under mild wash conditions. We apply PRISM to quantify the co-expression levels and nanometer-scale organization of one dozen cytoskeletal and synaptic proteins within individual neuronal synapses. Our approach is scalable to dozens of target proteins and is compatible with high-content screening platforms commonly used to interrogate phenotypic changes associated with genetic and drug perturbations in a variety of cell types.

## INTRODUCTION

Neuronal synapses are the fundamental sites of electrochemical signal transmission within the brain and are the primary cellular loci of plasticity that underlie learning and memory. Synapses are composed of dozens of proteins, whose expression levels, structural organization, and turnover govern diverse aspects of brain development and neuronal circuit function^1,2^. Because numerous synaptic protein genes have been implicated in psychiatric and neurological diseases^3–5^ and synaptic protein expression levels are known to vary widely across organisms, brain regions, and neuronal cell subtypes, characterizing synaptic protein composition and configuration in situ is of major importance to basic and translational neurobiology research. While fluorescence imaging offers the opportunity to characterize the heterogeneity in synaptic protein expression levels and localizations within intact neuronal samples^6^, it has been limited by its inability to visualize more than four protein species in any given neuronal sample using conventional imaging approaches.

Multiplexed imaging strategies that are used to overcome the spectral limit of conventional fluorescence microscopy typically involve multiple rounds of antibody staining and imaging achieved either by antibody elution^7,8^ or fluorophore inactivation using photo- and/or chemical bleaching^9–11^. For example, Array Tomography (AT) has been applied to volumetric imaging of synapses within intact brain tissue by sequentially staining and stripping the same ultrathin tissue sections with different antibodies^8,12,13^. More recently, gel embedding and expansion of whole intact organs has been used with sequential antibody loading and stripping to generate 13-channel fluorescence imaging datasets^14^. In addition, Cyclic Immunofluorescence (CycIF) has been applied to cancer cell lines to generate 9-channel diffraction-limited imaging using repetitive antibody loading-bleaching cycles. In each case, multiple antibody staining rounds were used together with harsh and time-consuming washing steps between imaging cycles, which may limit both epitope accessibility compared with simultaneous antibody loading, as well as alter epitope reactivity due to disruptive chemical or photobleaching treatment. Moreover, these preceding approaches are not readily amenable to super-resolution imaging within the same intact sample, and are therefore unable to resolve sub-synaptic protein structural organization. While electron microscopy (EM) has been incorporated into AT to facilitate correlative light and EM-based super-resolution imaging, EM is limited in its ability to resolve multiple molecular species in the same sample^15,16^, and requires complex sample fixation, embedding, and processing steps. In contrast to cyclic antibody staining-elution approaches, imaging mass cytometry (imaging CyTOF) and similar variants enable the detection of dozens of proteins within the same sample^17,18^. However, mass-spectrometry-based imaging requires sophisticated instrumentation that limits its broader utility, and its relatively low sensitivity compared with light microscopy renders reliable detection of synaptic proteins challenging.

In contrast, the use of diffusible, transiently binding fluorescent imaging probes that target specific neuronal protein targets in situ can, in principle, overcome each of the preceding limitations by offering, (1) simultaneous antibody loading prior to imaging; (2) multiplexing with gentle and rapid probe-exchange steps using only mild buffer treatment, in addition to (3) super-resolution imaging using PAINT (Points Accumulation In Nanoscale Topography)^19^. Originally introduced by Sharonov and Hochstrasser^19^, PAINT was first used to image reconstituted lipid membranes with nanoscale resolution using transient binding of diffusible dye molecules followed by probe localization and super-resolution image reconstruction. Subsequently, several variants of this approach^19–21^ were introduced, including uPAINT^20^ that employs diffusible fluorescent antibodies and DNA-PAINT^22,23^ that uses diffusible fluorescent single-stranded DNA (ssDNA) molecules (imaging probes) that transiently bind to complementary ssDNA oligos (docking strands) attached to target DNA nanostructures or antibodies to generate 10- or 4-channel data^23^, respectively. Protein-fragment-based probes have alternatively been used to generate multiplexed cytoskeletal and focal adhesion super-resolution images with higher labeling density compared with antibody-based approaches^21^. However, this strategy requires identification of highly specific, transiently binding peptides for each target molecular species, which may be challenging to generalize to other proteins, particularly those with lower expression levels than cytoskeletal proteins.

For each of the preceding super-resolution methods, time-lapse imaging is applied together with fluorophore localization and reconstruction algorithms commonly employed in the conventional photo-activation-based super-resolution imaging approaches Photoactivated Localized Microscopy (PALM)^24^ and Stochastic Optical Reconstruction Microscopy (STORM)^25^. Super-resolution imaging with ~20nm spatial resolution has enabled the visualization of sub-synaptic structures with fluorescent probes^26–28^, which is essential for the analysis of synaptic structures that range from 200 nm to 1 μm^26,27,29–32^. Average spatial distributions of synaptic proteins have been determined indirectly using three-channel 3D STORM by sequential imaging of distinct subsets of synaptic proteins in different samples, while keeping a common target in each round of imaging as a reference^26^. Despite their significantly enhanced resolution, however, these approaches are still subject to the conventional spectral limit of four distinct protein species that can be measured in a given sample. Imaging with transiently binding probes in principle enables unlimited multiplexing under physiological buffer conditions, and renders it feasible to apply both diffraction-limited and super-resolution imaging within the same, intact fixed sample by modulating probe affinity while retaining probe-binding specificity. Multiplexed imaging using diffusible imaging probes is also compatible with conventional commercial microscopes and high-content imaging systems, since the spectral limit is overcome through the use of probe exchange or washout, facilitating imaging-based phenotypic screens of genomic and drug perturbations.

Toward this end, here we introduce PRobe-based Imaging for Sequential Multiplexing (PRISM) that, unlike DNA-PAINT, utilizes either fluorescently labeled ssLNA or conventional ssDNA oligos as imaging probes to realize dual-purpose, multiplexed diffraction-limited confocal (LNA-PRISM) or PAINT-based super-resolution imaging using the same ssDNA-labeled detection antibodies or peptides (DNA-PRISM). Conventional DNA-PAINT cannot be applied for this purpose because of the high concentrations of low-affinity ssDNA imaging probes needed for confocal imaging, which increases fluorescence background and thus reduces the signal-to-background ratio of the images. In contrast, the use of ssLNA probes offers high-affinity binding that significantly reduces background fluorescence of the unbound, diffusible imaging probe species that makes confocal, diffraction-limited imaging possible. We apply LNA-PRISM to generate 13-channel confocal imaging data of 7 synaptic proteins imaged simultaneously with 5 cytoskeleton-related proteins, including filamentous actin and dendritic microtubules, in primary neuronal cultures imaged on a high-throughput confocal microscopy system in multi-well plate format. We additionally apply DNA-PRISM using the same ssDNA-antibody and -peptide conjugates to resolve the 20 nm-scale structural organization of 8 synaptic proteins together with filamentous actin and dendritic microtubules in situ. Our multiplexed confocal imaging data set enabled us to analyze 66 protein co-expression profiles extracted from thousands of individual synapses within the same intact neuronal culture, revealing strong correlations amongst subsets of synaptic proteins, as well as capturing heterogeneity in synapse sub-types. Our super-resolution imaging data revealed the nanometer-scale structural organization of 9 targets within individual synapses that is consistent with EM^33^ and average synaptic structure previously assayed using STORM^26^.

## RESULTS

### Overview of LNA- and DNA-PRISM

Our PRISM workflow employs neuronal cultures that are fixed, permeabilized, and stained simultaneously using ssDNA-conjugated antibodies or peptides, collectively termed "markers", which specifically label cellular targets. Markers are barcoded with single-stranded nucleic acid oligonucleotides ("docking strands”), rationally designed to maximize orthogonality between complementary fluorescently labeled ssLNA or DNA imaging probes used for confocal or super-resolution imaging (Figure 1a). To maximize the multiplexing capacity of the assay, primary rather than secondary antibodies are labeled with docking strands whenever possible so that labeling of distinct targets is not limited by the number of secondary antibodies reacting with different species. Extensive validation of each marker and fluorescently labeled ssLNA/ssDNA imaging probe is performed to ensure that markers retain their target-specific recognition properties following conjugation with nucleic acid docking strands, and that imaging strands target cognate docking strands with high affinity and specificity without cross-talk (Figure 1b). Following marker and imaging probe validation, multiplexed imaging is performed using sequential labeling and washing out of individual imaging probes, with wash-steps in between used to clear the sample of imaging probes (Figure 1c). Diffraction-limited, confocal images are acquired using LNA-PRISM, whereas single molecule time-lapse imaging followed by image reconstruction, drift correction, and image alignment is performed with DNA-PRISM using PAINT.

**Figure 1.**
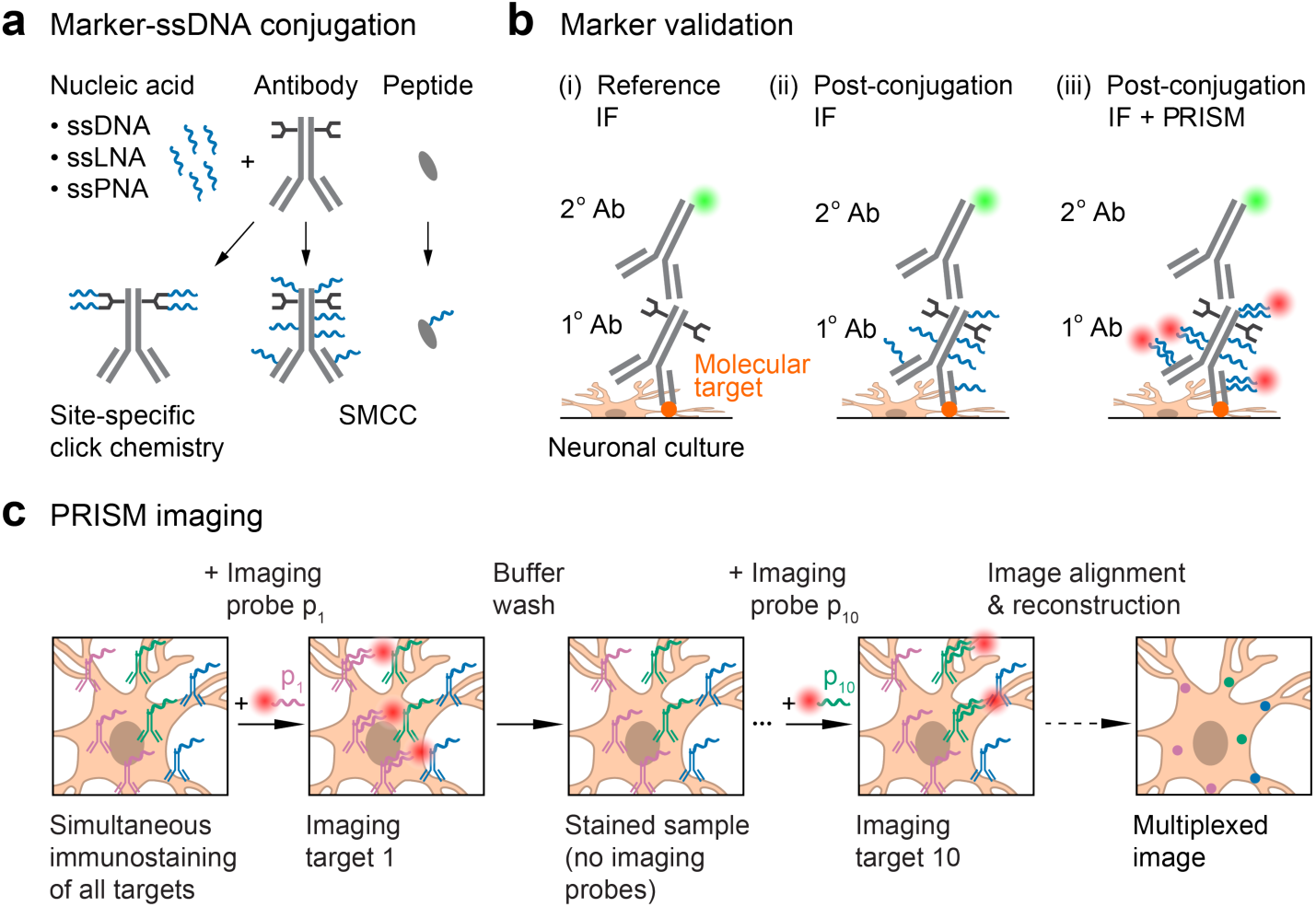
Schematic of the PRISM framework for highly-multiplexed imaging of molecular targets in neurons. (a) Reagents (“markers”, shown in gray) for detecting subcellular targets include antibodies or peptides that are conjugated with unique oligonucleotide barcodes (“docking strands”, shown in blue). A barcoded marker is imaged using the complementary fluorophore-conjugated oligonucleotides (imaging probes) that bind to the docking strands on the marker (see (iii) in (b), fluorophores shown as red circles). Binding affinity of the imaging probes to the docking strands can be varied by changing the sequence and type of the oligonucleotides, which thereby enables either diffraction-limited (high affinity) or superresolution microscopy (low affinity). Conjugation of docking strands to markers using site-specific click chemistry enables stoichiometric control of the number of nucleic acids bound to a whole antibody, while SMCC enables conjugation of docking strands to free amine groups on a variety of markers. (b) The reagent testing and validation phase consists of: (i) generating reference staining patterns of all molecular targets of interest using standard immunofluorescence (IF), (ii) Specificity and staining quality of markers conjugated with docking strands compared to those in the reference IF, and (iii) co-localization of PRISM-imaged staining patterns using imaging probes (red circles, which correspond to fluorophores conjugated to the probes) with standard IF staining patterns (green circles). (c) Overview of the main steps in the PRISM imaging workflow. All molecular targets of interest are immunostained at once using docking strand-conjugated markers (e.g., antibodies shown in green, blue, and pink). Nucleic acid imaging probes specific to each marker (e.g., p_1_−p_10_) are applied and imaged sequentially, with each imaging strand washed out after image acquisition at each step. This approach enables imaging a dozen or more distinct molecular targets in the same sample. Images of different markers are drift-corrected and overlaid to generate a pseudo-colored, multiplexed image. For super-resolution PRISM, prior to drift correction, the super-resolved image of each marker is reconstructed from the temporal image stack of binding/unbinding events of the imaging probes to/from the docking strands on the marker.

### Design and validation of markers for PRISM

To apply multiplexed neuronal imaging in either confocal or super-resolution modes using the same markers, we conjugated ssDNA docking strands to markers using either SMCC linkers or site-specific chemoenzymatic labeling (Materials and Methods). Whereas SMCC non-specifically conjugates docking strands to accessible primary amines on the marker through NHS chemistry, site-specific labeling conjugates ssDNA docking strands to four conserved glycan chains on the Fc region of the antibody (**Figure S1** and **Figure S2**), thereby minimizing the likelihood of disrupting the antibody paratope.

Because conjugation of antibodies and peptides with ssDNA may alter their affinity and/or specificity, we validated each marker-docking-strand conjugate individually in neuronal culture using indirect immunofluorescence (IF) to ensure the same staining patterns were obtained compared with the reference, unconjugated marker. However, most markers were found to exhibit strong nuclear localization following SMCC or site-specific conjugation with ssDNA, suggesting that the observed change in affinity of the ssDNA-conjugated antibody to its target is not solely due to the possible modification of paratopes by ssDNA (Figure 2a, **Figure S3** and **Figure S4**). To eliminate off-target nuclear localization of the ssDNA-conjugated antibodies, we screened several nuclear blocking agents and found that salmon sperm DNA successfully blocked the nuclear localization of ssDNA-conjugated antibodies (Figure 2a and **Figure S5**). Interestingly, conjugating antibodies with single-stranded Peptide Nucleic Acid (ssPNA)^34^ docking strands instead of ssDNA also eliminated nuclear localization in the absence of any blocking (Figure 2a), lending credence to the hypothesis that overall charge of the nucleic acid docking strands present on antibodies was responsible for their non-specific nuclear localization. Nevertheless, salmon sperm blocking was performed in all experiments to minimize cost and complexity associated with generating a full library of ssPNA docking strands. Image cross-correlation analysis showed that ssDNA-conjugated antibodies produced staining patterns similar to those of unmodified antibodies, as assessed through indirect IF when samples were blocked with salmon sperm DNA prior to immunostaining (Figure 2b and **Figure S6**).

**Figure 2.**
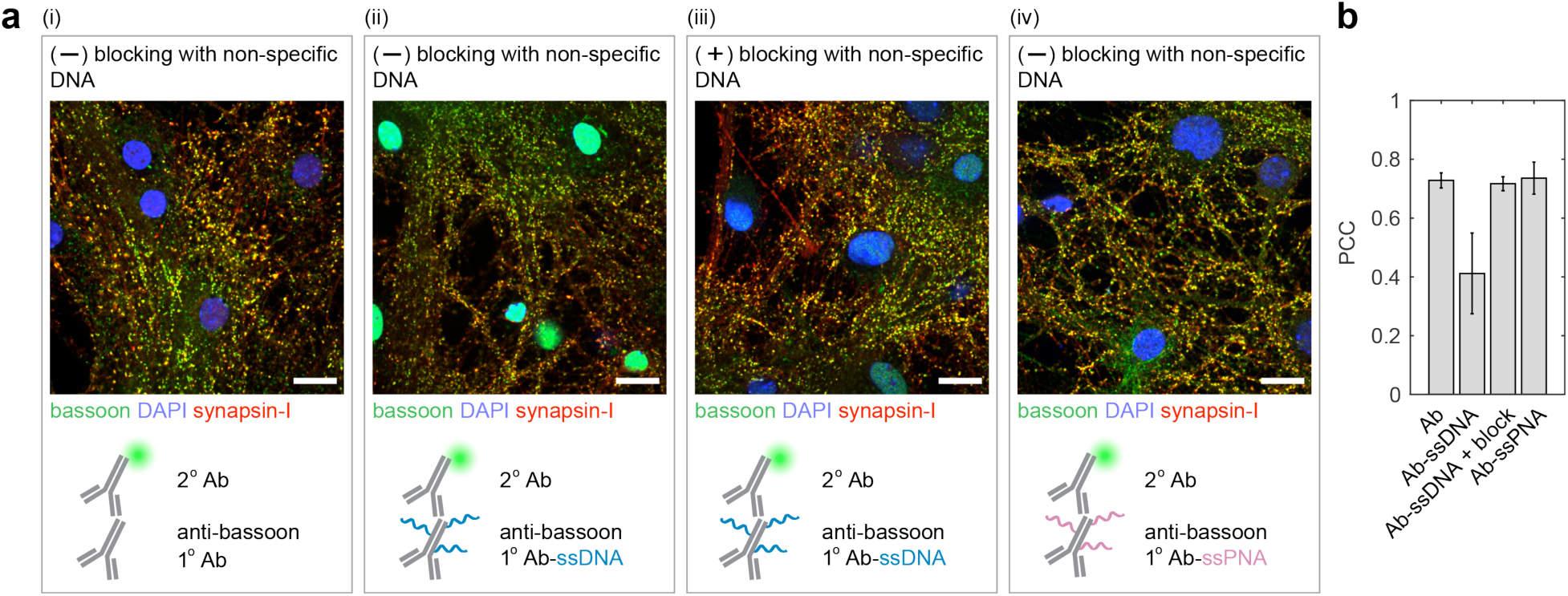
Blocking off-target nuclear localization of ssDNA-conjugated antibodies. (a) Neurons were stained with native or ssDNA-conjugated anti-bassoon antibody, anti-synapsin-I antibody, and DAPI. ssDNA-conjugated anti-bassoon antibody exhibited strong off-target nuclear localization (ii, green staining inside the nuclei) compared to the native antibody (i). This nuclear localization was reduced by blocking the fixed sample with non-specific (salmon sperm) DNA prior to immunostaining (iii), or when the anti-bassoon antibody used for staining was conjugated with ssPNA instead of ssDNA (iv). Scale bar, 20 μm. (b) Cross-correlation analysis of the IF images in (a). Pearson correlation coefficient (PCC) of the bassoon channel (green in (a)) with the synapsin-I channel (red in (a)) for each image. Differences in PCC indicate changes in antibody staining patterns. Error bars represent 95% confidence intervals.

The following strategy was employed when conjugating a new primary antibody: SMCC chemistry was first used to obtain high docking strand-to-marker labeling density for maximal PRISM signal. For antibodies that changed their localization pattern after SMCC conjugation and could not be rescued using nuclear blocking, site-specific antibody conjugation was used, or, alternatively, a different primary antibody was tried instead, granted that another high-quality primary antibody was available for the target of interest. If staining patterns still changed with site-specific conjugation or with a different primary antibody, ssDNA-conjugated secondary antibodies were employed to facilitate visualization of the target.

### Imaging probe design for LNA-PRISM

To enable high-throughput confocal imaging of neurons with high signal-to-noise and low background fluorescence in multi-well plate format, we designed high affinity ssLNA imaging strands of 11 nt length that target the same 11 nt ssDNA docking strands used in DNA-PAINT imaging with high specificity^23^. Similar to ssDNA, ssLNA binding affinity to complementary ssDNA is salt-dependent, thereby enabling rapid probe exchange via imaging probe wash-out using low salt concentration buffer (**Figure S7**). Orthogonality of each imaging probe was validated individually using a cell-based crosstalk assay that resembles the staining and imaging conditions in a multiplexed PRISM experiment. Results of this cross-talk assay showed <10% crosstalk between each of the docking-imaging-strand pairs for our staining and imaging conditions (**Figure S8** and **Figure S9**). Typical imaging strand incubation and wash-out times for LNA-PRISM are 5–10 minutes each, which is considerably faster than existing multiplexed imaging approaches that require multiple rounds of antibody staining and elution that can require up to hours or days to complete^8,10,12^. PRISM washing conditions consist of 0.01× phosphate buffered saline (PBS), which is also milder than alternative multiplexed imaging methods that utilize oxidizing reagents or high-pH buffer^8,10,12^. In addition to reducing the risk of altering epitopes over the course of multiple wash cycles, mild buffer conditions minimize the possibility of the disruption of delicate cellular structures, which may be particularly crucial for preserving the integrity of neuronal synapses. RNase was used to eliminate off-target binding of ssLNA imaging probes to cellular RNA (**Figure S10**), a treatment that did not affect antibody marker localization (**Figure S11**). RNase-treated cells produced target staining patterns using LNA-PRISM that were indistinguishable from those with conventional indirect IF (**Figure S12**).

### LNA-PRISM: 13-channel confocal neuronal imaging

13-plex imaging of cultured rat hippocampal neurons using 10 ssLNA imaging probes and three non-PRISM fluorescent markers was performed to characterize the synaptic and cytoskeletal protein-protein network that is core to the regulation of synapse formation and plasticity (Figure 3a). This network includes the cytoskeletal proteins actin, Tuj-1, MAP2, ARPC2, and cortactin, the pre-synaptic proteins synapsin-I, bassoon, and VGLUT1, the post-synaptic density proteins PSD-95, Homer-1b/c, and SHANK3, and the receptor NMDAR2B. The canonical synaptic markers synapsin-I, bassoon, VGLUT1, PSD-95, Homer-1b/c, and SHANK3 exhibited a high degree of co-localization, with punctate patterns, whereas Tuj-1 and MAP2 yielded clear cytoskeletal morphologies (Figure 3b). Noticeably, ARPC2 and cortactin displayed punctate patterns that also co-localized with other synaptic markers, in agreement with previous results^35^. To assess whether multiple rounds of imaging probe wash-out and probe application steps physically distorted the sample or noticeably stripped markers from their epitopes, synapsin-I was imaged twice, once in the middle and once at the end of the PRISM experiment, which revealed highly reproducible localization patterns (Figure 3a).

**Figure 3.**
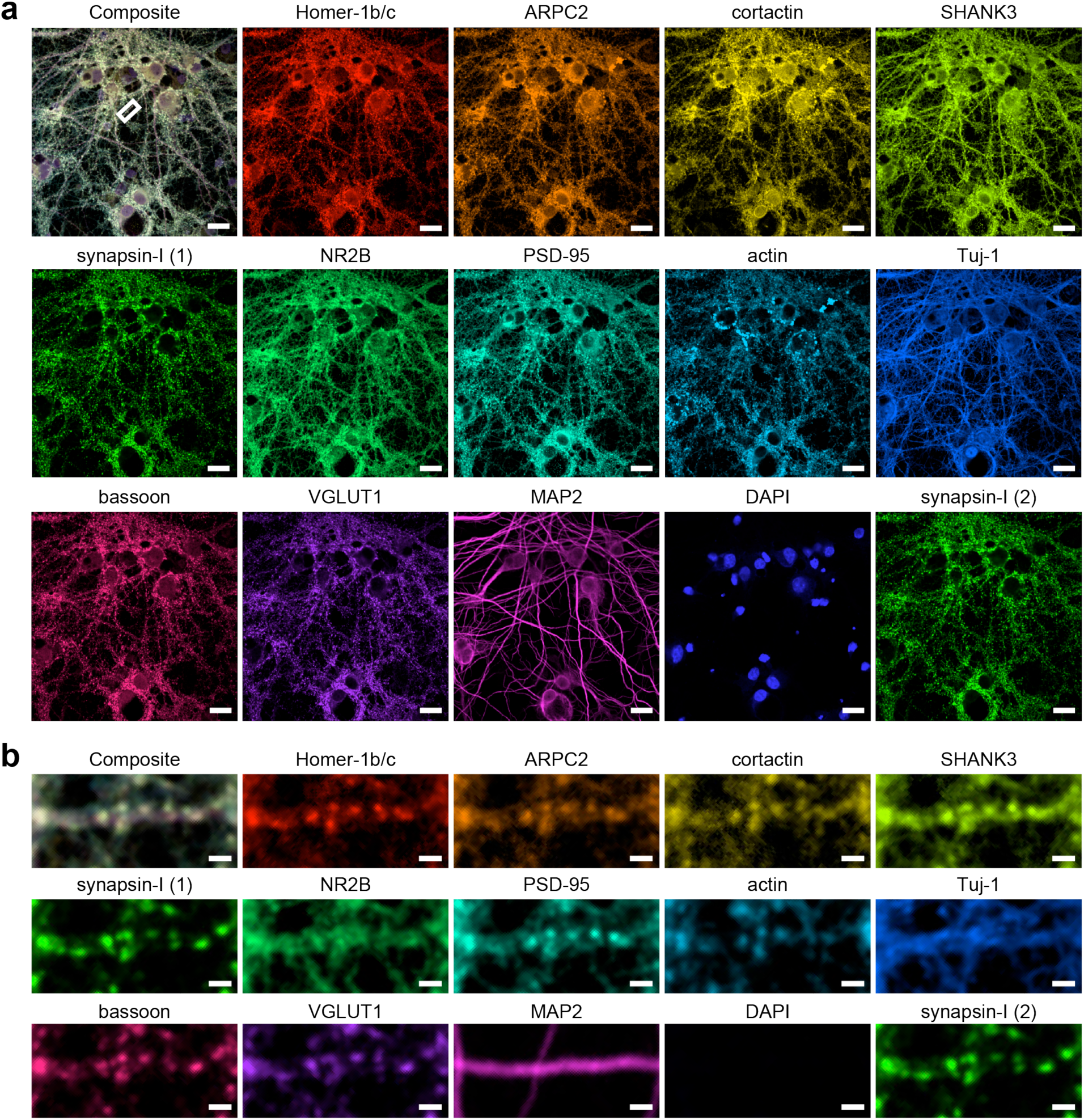
| Confocal LNA-PRISM image of rat hippocampal neuronal synapses. (a) 13-channel images of 21 days in vitro (DIV) rat hippocampal neuronal culture. The overlaid image is shown in the top-left corner, followed by the image of each individual channel. DAPI, MAP2 and VGLUT1 were visualized using fluorescently labeled secondary antibodies, while other targets were visualized using ssLNA imaging probes. Synapsin-I was imaged twice, once in the middle and once at the end of the experiment. (b) Zoom-in view of a single dendrite indicated by the white box in (a). Scale bars: (a) 20 μm; (b) 2 pm.

LNA-PRISM offers nearly an order of magnitude increase in the ability to detect co-localization patterns in situ, with 66 protein-protein co-localizations using 12 protein labels compared with only 6 from conventional 4-channel imaging. Individual synaptic features including size and intensity were extracted for each target from PRISM images using an image-processing pipeline optimized for synapse segmentation (**Figure S13**, Materials and Methods). This analysis established correlations between distinct synaptic features computed across all synapses, with a high correlation score between two synaptic proteins indicating a potential functional association. Expression levels of most synaptic proteins were highly correlated, with the exception of the cytoskeletal proteins Tuj-1 and MAP2, in agreement with previous image cross-correlation analyses that showed that tubulin is largely excluded from synapses^12^ (Figure 4a). In addition, the post-synaptic density proteins Homer-1b/c, PSD-95, and SHANK3 strongly correlated with one another in their localization patterns, which may be attributed to their dense and compact protein distributions within the PSD^36,37^. The Arp2/3 complex subunit ARPC2, which has been shown to interact with SHANK3 within synapses^35^, also correlated in expression level with other PSD proteins. In agreement with the expected separation of pre-and post-synaptic localization sites, proteins such as synapsin-I and VGLUT1 that are associated with synaptic vesicles were highly correlated with the pre-synaptic scaffolding protein bassoon, but were only weakly correlated with the post-synaptic proteins (Figure 4a). Interestingly, bassoon exhibited moderate correlation in expression with all of the PSD proteins, as shown previously in the mouse olfactory bulbs^26^. These data suggest a coordination of pre- and post-synaptic structures across the synaptic cleft, which may be used in future knock-down or other screens to discover novel trans-synaptic protein-protein associations.

**Figure 4.**
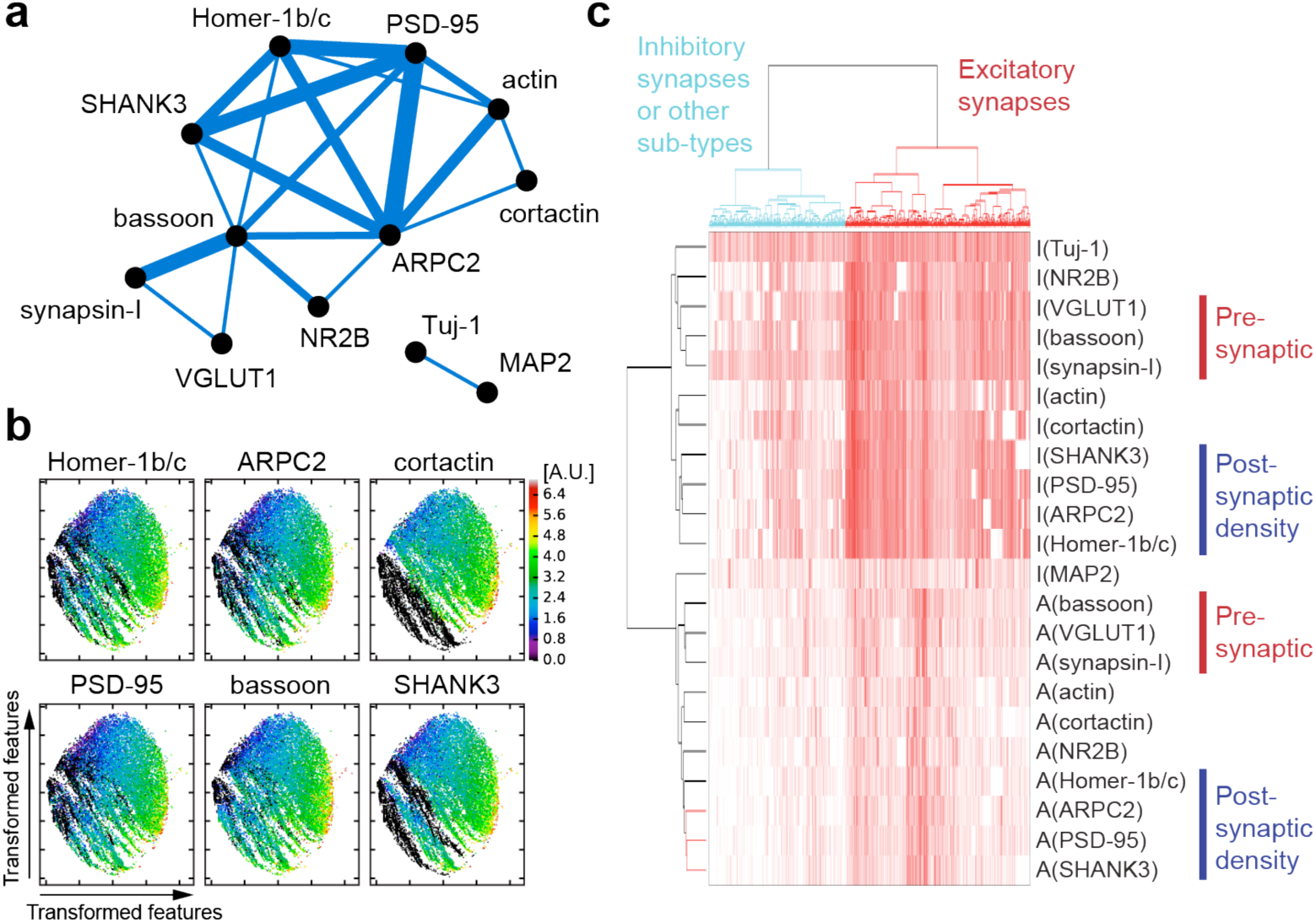
| Analysis of single-synapse profiles from multiplexed confocal imaging data acquired using LNA-PRISM. (a) Network representation of correlations between intensity levels of synaptic proteins within synapses (n=175,399 synapses from 3 cell culture batches). The thickness of each edge represents the relative correlation strength between the respective nodes. (b) t-Distributed Stochastic Neighbor Embedding (t-SNE) maps of n=17,894 synapses from a single culture batch; each with 20 features (intensity levels and punctae sizes of 10 synaptic proteins). Each point in each t-SNE map represents a single synapse with its (x,y) coordinates corresponding to the transformed features that best preserve the distribution of synapses in the original high dimensional feature space. Intensity levels of individual proteins are color-coded in each map. (c) Hierarchical clustering analysis of synapse profiles. Each column in the heat map represents a profile of a single synapse with 24 synaptic features (rows). “I” and “A” denote image intensity level and punctum size, respectively (n=53,682 synapses from a single culture batch).

Large-scale profiling of synapses also enabled us to use the rich protein co-expression feature profiles assayed with LNA-PRISM to classify synapse subtypes. To identify putative subcategories of synapse types in this high-dimensional feature space that consists of 20 dimensions (intensity levels and punctae sizes for 10 synaptic proteins), we applied t-Distributed Stochastic Neighbor Embedding (t-SNE), a tool commonly used to visualize high-dimensional single-cell data (Figure 4b)^38^. t-SNE transforms high dimensional data into two dimensions, aiming to preserve the local high dimensional data structure within the lower dimensional space. t-SNE analysis of 17,894 synaptic profiles revealed that the majority of synapses contain most of the synaptic proteins that we measured, which, given that our antibody panel consisted mostly of excitatory proteins, also likely corresponded to excitatory synapses. In addition, smaller sub-type clusters were identified, showing an absence of one or more synaptic proteins, which may correspond to additional synapse subtypes. Hierarchical clustering of protein feature profiles corroborated findings of the preceding correlation and t-SNE analyses, namely that pre-synaptic proteins are highly clustered with one another, whereas PSD proteins and ARPC2 form a separate sub-cluster (Figure 4c). These findings suggest that protein associations derived from PRISM data recapitulate the molecular composition and structural properties of excitatory synapses, and can do so for one dozen targets simultaneously in thousands of synapses within the same intact sample imaged within hours in multi-well plate format.

### DNA-PRISM: Super-resolution imaging using low affinity ssDNA imaging strands and PAINT

The same antibody-ssDNA conjugates offered the ability to also super-resolve molecular targets within individual synapses in primary mouse neuronal cultures using DNA-PRISM (DNA-PAINT)^22,23^. Neuronal cultures were assembled into flow cells in which fluid exchange was controlled by an automated fluidics handling system to ensure gentle buffer washing and imaging probe application designed to minimize sample distortion. Super-resolution DNA-PRISM images of microtubules and F-actin in neurons were first compared with widefield IF images. Super-resolved microtubules formed bundles within neuronal processes, whereas f-actin exhibited linear, filamentous structures within these regions, but showed punctae within dendritic spines (Figure 5a-d). Subcellular structures imaged with DNA-PRISM correlated well with the corresponding widefield images, but with significantly improved spatial resolution (Figure 5b,d). A Gaussian fit to the cross-sectional profile of a single microtubule produced a full-width at half-maximum (FWHM) of 46.5 nm (Figure 5g), which is consistent with previous PAINT measurements in HeLa cells^23^. In addition to microtubules and f-actin, DNA-PRISM imaging of neuronal synapses also corresponded well with the widefield IF images, but with significantly improved resolution, as expected, revealing closely apposed pre- and post-synaptic sites (Figure 5e-f). We quantified the average synapse size defined by synapsin-I and PSD-95 punctae using the radial cross-correlation function between synapsin-I and PSD-95, with the decay length of the correlation function revealing an average synapse size of ~200 nm (Figure 5i)^12^. The spatial decay of the DNA-PRISM correlation curve occurred at a smaller spatial scale than the widefield imaging curve, indicating the smaller synapse size revealed by DNA-PRISM due to enhanced resolution relative to widefield imaging.

**Figure 5.**
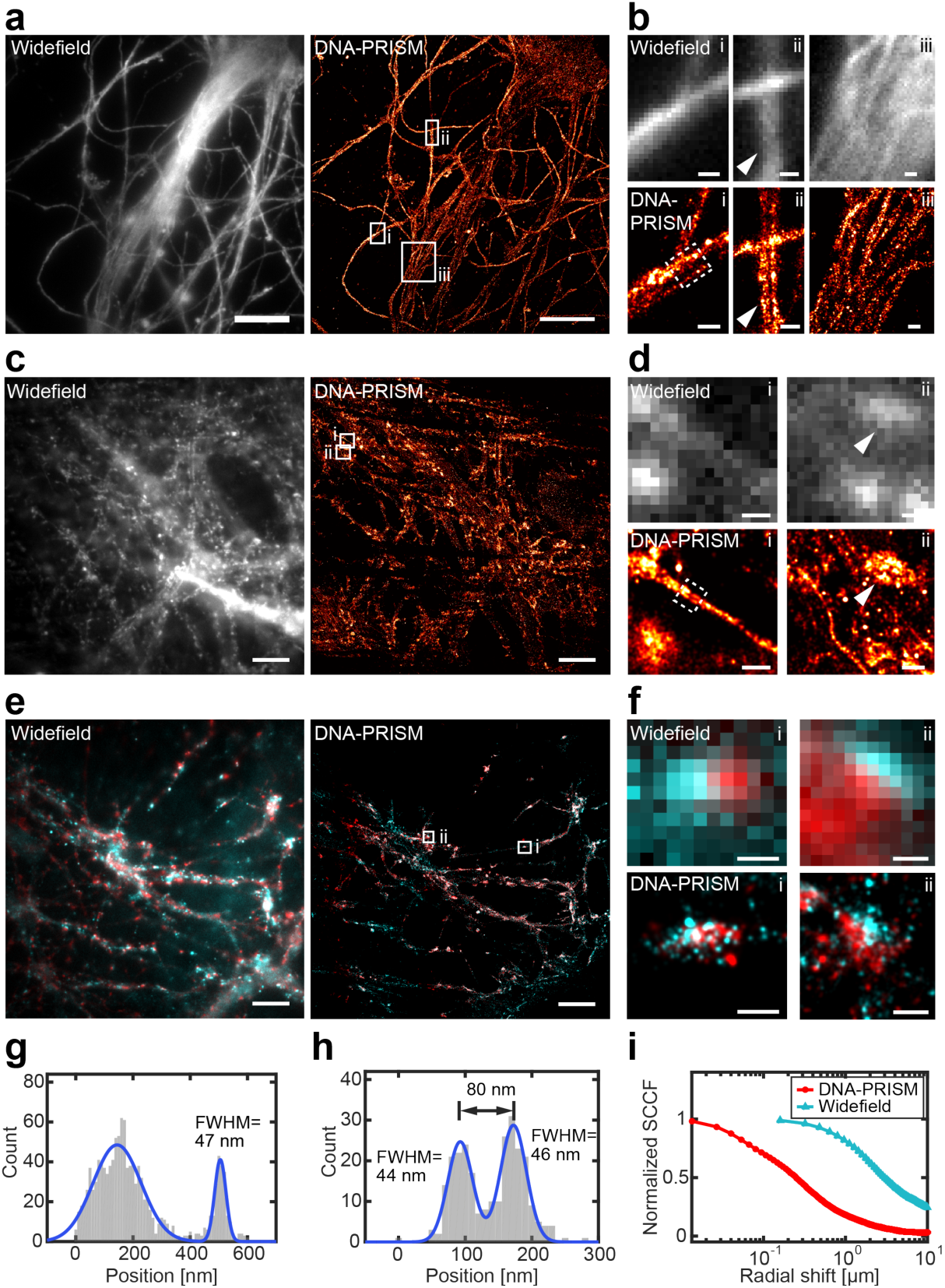
Super-resolution DNA-PRISM imaging of primary neuronal cultures. (a) Widefield and DNA-PRISM images of neuronal microtubules stained using the DNA-conjugated anti-Tuj-1 antibody. (b) Zoom-in view of the boxed areas in (a) show resolution enhancement of DNA-PRISM images compared with widefield images. The arrowhead indicates distinct microtubule bundles that are not resolved in the widefield image (c) Widefield and DNA-PRISM images of filamentous actin stained using DNA-conjugated phalloidin. (d) Zoom-in views of the boxed areas in (c) show two actin filaments (left) and the synaptic actin punctae with sub-synaptic structures (right, arrow head) that are not resolved in widefield images. (e) Widefield and DNA-PRISM images of pre-synaptic marker synapsin-I (red) and post-synaptic marker PSD-95 (cyan) of the same field of view. (f) Zoom-in view of single synapses indicated by boxes in (e). (g) Cross-sectional profile of the boxed region in (b) shows a microtubule bundle next to a possible single microtubule with FWHM = 47 nm. (h) Cross-sectional profile of the boxed region in (d) shows two actin filaments or small filament bundles that are 80 nm apart. (i) The average size of synapses defined by synapsin-I and PSD-95 is quantified using the normalized radial crosscorrelation function. The decay at the smaller radial shift of the DNA-PRISM curve (red) indicates the smaller synapse size in the DNA-PRISM image due to the improved spatial resolution. Scale bar: (a,c,e) 10 μm; (b,d,f) 0.5 μm.

We next applied DNA-PRISM imaging to super-resolve the nanoscale pre- and post-synaptic organization of 9 targets within individual synapses, including Tuj-1, f-actin, cortactin, PSD-95, synapsin-I, NMDAR2B, SHANK3, Homer-1b/c, and bassoon (Figure 6a). Due to differences in synapse orientations with respect to the imaging plane, individual synapses varied in their degree of overlap among proteins within each synapse. For a subset of synapses with the proper orientation relative to the imaging plane, we identified clear separation between pre-synaptic proteins (synapsin-I and bassoon) and post-synaptic proteins (PSD-95, SHANK3, Homer-1b/c)(Figure 6b). Pre- and post-synaptic proteins were localized on opposite sides of the synaptic cleft, whereas cytoskeletal proteins (Tuj-1, actin, cortactin) were observed at both sides of the cleft. Moreover, PSD proteins (PSD-95, SHANK3, Homer-1b/c) showed narrow, overlapping distributions in expression levels, suggesting physical interaction of these proteins in the PSD (Figure 6c). In contrast, synapsin-I exhibited a broader spatial distribution compared with the distributions of scaffolding proteins, in agreement with the more diffuse distributions expected for vesicle-associated proteins (Figure 6c). These spatial distributions of synaptic proteins were consistent with the average distributions previously measured from multiple synapses and distinct cultures using three-channel STORM^26^ and EM^33^. However, in stark contrast to these previous studies that relied on reference markers, our imaging and analysis of sub-synaptic proteins resolved all measured targets of interest within the same synapse simultaneously. Integration of our approach with 3D super-resolution imaging systems would offer its application to dozens or hundreds of synapses in situ.

**Figure 6.**
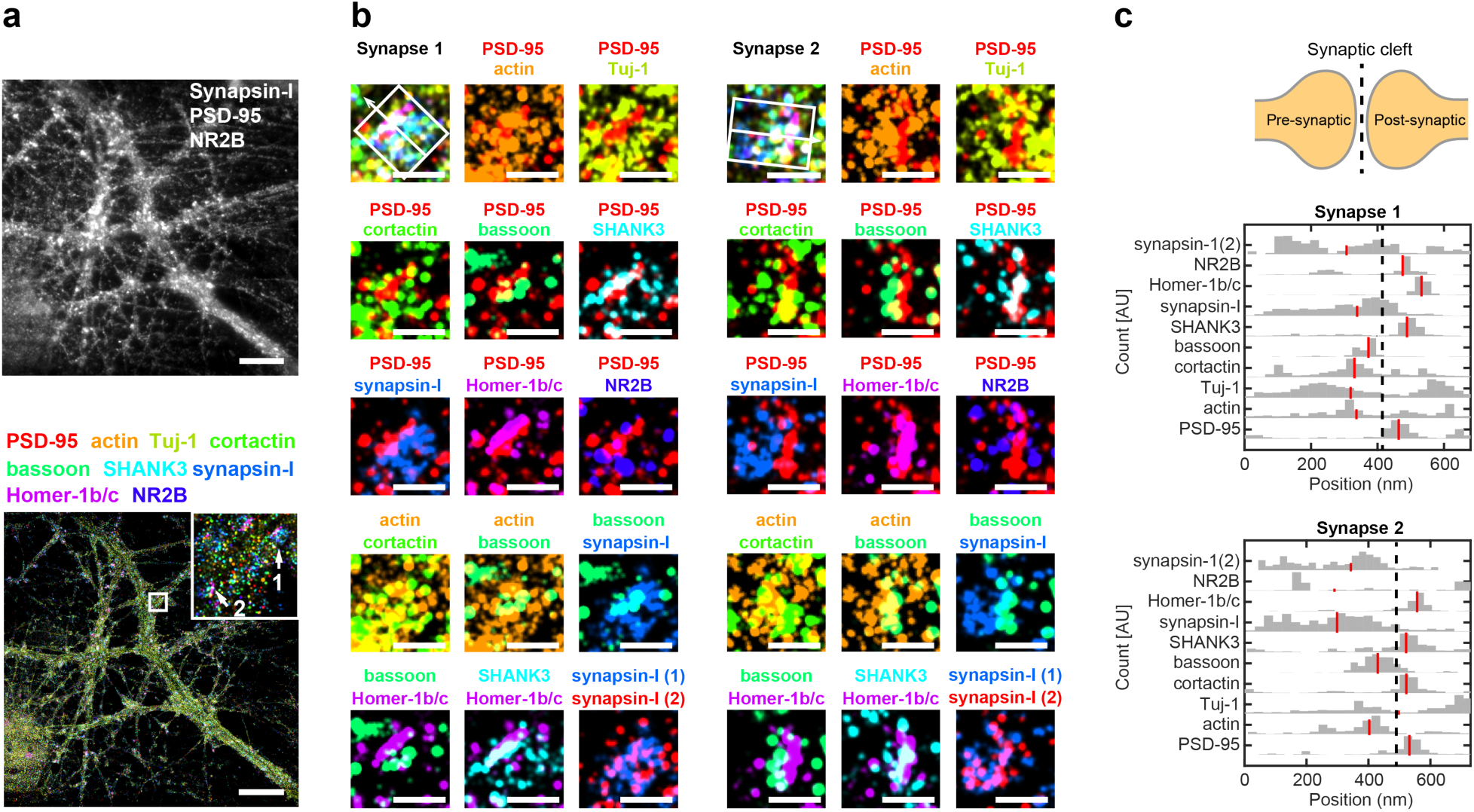
Multiplexed DNA-PRISM image shows distributions of synaptic proteins within individual synapses. (a) Single-channel widefield image of three targets (top) and nine-channel DNA-PRISM images (bottom) of the same field of view. (b) Zoom-in view of two individual synapses in (a) shows the separation of pre-synaptic proteins (synapsin-I and bassoon) and post-synaptic proteins (PSD-95, SHANK3, Homer-1b/c). For each synapse, the nine-target image is shown in the top-left corner, with distinct pairs of synaptic proteins shown in the remaining images. Synapsin-I was imaged twice, once at the beginning and once at the end of the experiment. (c) Cross-sectional profiles of protein distributions along trans-synaptic axes (white boxes with arrows in (b)) of the two synapses in (b). Red lines indicate the medians of the distributions.

## DISCUSSION

PRISM offers a powerful and versatile approach to multiplexed fluorescence imaging, phenotypic profiling, and high-resolution structural analysis of fixed neuronal cultures. In one variant, LNA-PRISM utilizes high-affinity ssLNA imaging probes to realize high-throughput confocal imaging for rapid, large-scale screening of neuronal phenotype applied here to over one-dozen cytoskeletal and synaptic protein targets including tens of thousands of individual synapses in triplicate. In a second variant, DNA-PRISM utilizes the same antibody/peptide reagents with ssDNA imaging probes to perform super-resolution synaptic imaging with PAINT. In future studies, large-scale morphological screens may first be performed in multi-well plate format using LNA-PRISM, followed by super-resolution imaging of a sub-set of synapses or neuronal subregions with DNA-PRISM to resolve synaptic ultra-structure. Compared with previous multiplexed diffraction-limited imaging approaches that utilize sequential antibody labeling and stripping or bleaching^8,10,12^, LNA-PRISM offers simultaneous staining of all protein targets, which reduces the risk of "masking” antigens, as well as substantially increasing assay throughput. The use of physiological wash buffers additionally minimizes the possibility of sample or epitope disruption, which may be crucial for high-resolution structural and co-localization studies requiring high sample fidelity, such as in the profiling of synapses within cultured neuronal samples.

Application of our primary antibody conjugation strategy together with the use of libraries of orthogonal ssDNA sequences^39^ offers the potential for future application of PRISM to neuronal and other cellular systems exceeding substantially the approximately one dozen targets realized here. Moreover, genetic and drug perturbation screens aimed at discovering subtle alterations in neuronal phenotype should benefit substantially from the large-scale protein association networks within synapses that are derived from the 12 synaptic targets examined in this study, which offer 66 pair-wise synaptic co-localizations within the same neuronal culture. While single-color imaging probes were used to demonstrate the robustness of our probe-exchange strategy, which yielded reproducible protein localization even after 10 successive probe exchanges, future applications that utilize multiple laser lines to simultaneously image 3 distinct fluorophores simultaneously in any given imaging cycle renders our approach viable for multiplexing at least 30 molecular targets in situ. The significant increase in phenotypic information content realized by our approach offers major potential for both basic and translational neuroscience research, including high-content screening of phenotypic variation due to genetic and compound perturbations, as well as super-resolution ultra-structural synaptic imaging with nanometer-scale resolution. Future application of PRISM to light-sheet and 3D super-resolution imaging may also enable multiplexed analysis of neuronal morphology and nanoscale protein organization within fixed human and diverse model organism tissues and organoids.

## MATERIALS AND METHODS

### SMCC (succinimidyl 4-(N-maleimidomethyl)cylohexane-1-carboxylate) ssDNA conjugation of antibodies and phalloidin

Twenty five nanomoles of thiolated single-stranded DNA (ssDNA)(Integrated DNA Technologies, Inc., Coralville, IA) was reduced using Dithiothreitol (DTT, 50 mM) for 2 h, purified using a NAP-5 column (GE Healthcare Life Sciences, Inc., Marlborough, MA), and quantified using a NanoDrop 2000 (Thermo Fisher Scientific, Inc., Waltham, MA). In parallel, 100 μg of antibody was concentrated to 1 mg/mL using an Amicon Ultra 0.5 mL centrifugal column (100 kDa, EMD Millipore, Inc., Billerica, MA) and purified from additive chemicals such as sodium azide using a Zeba spin desalting column (7 kDa, Thermo Fisher Scientific). A freshly prepared solution of SMCC with 5% DMF in PBS (Sigma-Aldrich, Inc., St. Louis, MO) was then added to react for 1.5 h with the antibody solution at a molar ratio of 7.5:1. Unreacted SMCC was removed using a 7 kDa Zeba column. In a subsequent reaction, antibodies were mixed in a 1:15 molar ratio with the reduced thiolated ssDNA strands and incubated overnight at 4 °C to form stable thioether bonds. ssDNA-conjugated antibodies were then purified using an Amicon Ultra 0.5 mL column (100 kDa). The final protein concentration of the antibody was measured using a Nanodrop 2000 and the antibodies were stored at −20 °C in 1× PBS with 50% glycerol. Amino-modified phalloidin (Bachem AG, Bubendorf, Switzerland) was conjugated using the protocol described above with an extra step of HPLC purification to separate unreacted thiolated ssDNA and ssDNA-conjugated peptides. See **Table S1** for antibody information.

### Site-specific ssDNA conjugation of antibodies

The SiteClick™ antibody labeling system (Thermo Fisher Scientific) enables site-specific conjugation to four conserved glycan sites present on the Fc region of the heavy chains using copper-free click chemistry. Briefly, 100 μg of each antibody was concentrated to 2−4 mg/mL in azide-free Tris buffer and treated with β-galactosidase enzyme to modify carbohydrate domains. In a second step, azide modified sugars were attached to the modified glycan chain using β-1,4-galactosyltransferase. After overnight incubation and purification of antibodies using an Amicon Ultra 0.5 mL centrifugal column (100 kDa), DBCO-modified ssDNA docking strands (Integrated DNA Technologies, Inc.) were mixed at a 40× molar ratio with azide-modified antibodies and incubated overnight at 25°C. ssDNA conjugated antibodies were then purified using an Amicon Ultra 0.5 mL (100 kDa). Final concentration was measured using a Nanodrop 2000 and antibodies were stored in 1× PBS (Sigma-Aldrich) with 50% glycerol. For PNA-antibody conjugation, DBCO-modified mini-PEG-Y-PNA strands^34^ (PNA Innovations, Inc., Woburn, MA) were used as the docking strands. See **Table S1** for antibody information.

### Fluorophore conjugation of ssLNA imaging probes

3’-amino ssLNA strands were purchased from Exiqon A/S and conjugated with ATTO 655-NHS (ATTO-TEC GmbH, Siegen, Germany). Briefly, 10 nmoles of ssLNA were mixed with ATTO 655-NHS in 1:5 molar ratio in 500 uL PBS (Sigma-Aldrich), incubated for 2 h at room temperature and then overnight at 4 °C. Prior to purification with HPLC, isopropanol precipitation was performed to remove free dyes: 50 uL of 3M sodium acetate solution was first added to the ssLNA solution followed by adding 550 uL of isopropanol (−20 °C). The mixed solution was thoroughly vortexed and incubated for 30 min at −20 °C, and then immediately centrifuged at 10,000 g and 4 °C for 1 h. The supernatant was delicately removed from the pellet. 500 uL of cold ethanol (−20 °C) was carefully added to the pellet and centrifuged at 10,000g and 4 °C for 15 min. The supernatant was removed, and the pellet was dried at 37 °C for 2 h. The pellet was then resuspended in 500 uL PBS (Sigma-Aldrich), purified with HPLC, and lyophilized. See **Table S2** for ssLNA probe sequences.

### SDS-PAGE and mass spectrometry validation of conjugated antibodies

ssDNA-modified antibody solutions were reduced with 10 mM Tris-HCl complemented with DTT (20 mM) for 2 h at 37 °C. Reduced antibody solutions were then run on an SDS-PAGE gel (10% acrylamide/bis-acrylamide), for 90 min at 110 V. Staining was performed using EZBlue^™^ gel staining reagent (Sigma-Aldrich). SiteClick^™^ conjugation efficiency and the ssDNA to antibody ratio (DAR) were determined using MALDI-TOF mass spectrometry. Briefly, 20 μL of modified antibody solutions in 1× PBS (Sigma-Aldrich) (0.5–1 mg/mL) were purified and concentrated using ZipTip^®^ pipette tips C_4_ resin (EMD Millipore) and then eluted in 10 μL of 80% ACN 0.1% TFA, dried down, and re-constituted in 1 μL of sinapinic acid matrix solution. The samples were then spotted and analyzed with microflex MALDI-TOF (Bruker Daltonics, Inc., Billerica, MA).

### Primary mouse and rat neuronal cultures

Procedures for mouse neuronal culture preparation were approved by the Massachusetts Institute of Technology Committee on Animal Care. Hippocampal and cortical mouse neuronal cultures were prepared from postnatal day 0 or day 1 Swiss Webster (Taconic, Inc., Germantown, NY) mice as previously described^40,41^ but with the following modifications: dissected hippocampal and cortical tissues were digested with 50 units of papain (Worthington Biochem, Inc., Lakewood, NJ) for 6–8 min, and the digestion was stopped with ovomucoid trypsin inhibitor (Worthington Biochem). Cells were plated at a density of 10,000 per well in a glass-bottom 96-well plate coated with 50 μl Matrigel (BD Biosciences, Inc., San Jose, CA). Neurons were seeded in 50 μl Plating Medium containing MEM (Life Technologies, Carlsbad, CA), glucose (33 mM, Sigma-Aldrich), transferrin (0.01%, Sigma-Aldrich), Hepes (10 mM), Glutagro (2 mM, Corning, Inc., Corning, NY), Insulin (0.13%, EMD Millipore), B27 supplement (2%, Thermo Fisher Scientific), heat inactivated fetal bovine serum (7.5%, Corning). After cell adhesion, additional Plating Medium was added. AraC (0.002 mM, Sigma-Aldrich) was added when glia density was 50–70%. Neurons were grown at 37 °C and 5% CO_2_ in a humidified atmosphere.

Procedures for rat neuronal culture were reviewed and approved for use by the Broad Institutional Animal Care and Use Committee. For rat hippocampal neuronal cultures, E18 embryos were collected from CO_2_ euthanized pregnant Sprague Dawley rats (Taconic). Embryo hippocampi were dissected in ice-cold Hibernate E supplemented with 2% B27 supplements and 100U/mL Penn/Strep (Thermo Fisher Scientific). Hippocampal tissues were digested in Hibernate E containing 20U/mL papain, 1mM L-cysteine, 0.5mM EDTA (Worthington Biochem) and 0.01% DNAse (Sigma-Aldrich) for 8min, and the digestion was stopped with 0.5% ovomucoid trypsin inhibitor (Worthington Biochem) and 0.5% bovine serum albumin (BSA)(Sigma-Aldrich). Neurons were dissociated and plated at a density of 15,000 cells/well onto poly-D-lysine coated, black-walled, thin-bottomed 96-well plates (Greiner Bio-One, Inc., Kremsmünster, Austria). Neurons were seeded and maintained in NbActiv1 (BrainBits, Inc., Springfield, IL). Cells were grown at 37 °C in a 95% air with 5% CO_2_ humidified incubator for 21 days before use. All procedures involving animals were in accordance with the US National Institutes of Health Guide for the Care and Use of Laboratory Animals.

### Immunostaining and analysis for validation of ssDNA-conjugated antibodies

To test whether the binding specificities of antibodies were affected by ssDNA conjugation, immunostaining patterns of unconjugated and ssDNA-conjugated antibodies were compared in each case. Cells were fixed at room temperature for 15 min with 4% paraformaldehyde (PFA) (Electron Microscopy Sciences, Inc., Hatfield, PA) and 4% wt/vol sucrose (Sigma-Aldrich) in PBS (Sigma-Aldrich), and then washed three times with PBS. Cells were permeabilized for 10 min at room temperature with 0.25% Triton-X100 in PBS and washed twice with PBS. For staining with unconjugated primary antibody, cells were blocked for 1 hr at room temperature with the regular blocking buffer (5% BSA (Sigma-Aldrich) in PBS). Cells were then incubated with primary antibodies diluted in the blocking buffer overnight at 4 °C. For staining with ssDNA-conjugated primary antibodies, the nuclear blocking buffer (5% BSA and 1 mg/mL salmon sperm DNA (Sigma-Aldrich) in PBS) was used instead of regular blocking buffer for blocking and antibody dilution. After primary antibody staining, the sample was then washed three times with PBS, incubated for 1 hr at room temperature with secondary antibodies in 5% BSA in PBS, and washed again three times with PBS. For validation of ssDNA-conjugated secondary antibodies, the fluorescently labeled secondary antibodies of the same species were added to the samples after 30 min incubation with ssDNA-conjugated secondary antibodies to reduce the competition of binding of fluorescently labeled secondary antibodies with ssDNA-conjugated secondary antibodies. See **Table S1** for antibody information. Comparison of antibody staining patterns before and after ssDNA conjugation were performed using the Pearson Correlation Coefficient (PCC). Colocalization of each antibody being tested with synapsin-I signal was performed before and after ssDNA conjugation. Specifically, three confocal images were acquired of neurons stained with unconjugated and conjugated antibodies separately. Each image was split into four quadrants, and the PCC between the synapsin-I channel and the channel of the other synaptic antibody for each quadrant was calculated and then averaged to obtain the mean PCC.

### Immunostaining for LNA- and DNA-PRISM

Cells were fixed and permeabilized as described in the previous section. For LNA-PRISM, cells were additionally incubated in RNase solution (50 μg/mL RNase A (Thermo Fisher Scientific) and 230 U/mL RNase T1 (Thermo Fisher Scientific) in 1× PBS (Sigma-Aldrich)) at 37 °C for 1 h to reduce the fluoresce background caused by ssLNA-RNA binding, and washed 3 times with PBS. Cells were then blocked for 1 hr at room temperature with the regular blocking buffer (5% BSA (Sigma-Aldrich) in PBS). The following unconjugated primary antibodies were diluted in the regular blocking buffer and used for LNA- or DNA-PRISM: MAP2, VGLUT1, PSD-95, and NMDAR2B (LNA-PRISM); PSD-95 and NMDAR2B (DNA-PRISM). Cells were incubated in diluted primary antibodies overnight at 4 °C, washed 3 times with PBS, and then incubated in the nuclear blocking buffer for 1 h at room temperature. Next, the following secondary antibodies were diluted in the nuclear blocking buffer and used for LNA- or DNA-PRISM: goat-anti-chicken-Alexa 488, goat-anti-guinea pig-Alexa555 and goat-anti-rabbit-p1, goat-anti-mouse-p12 (LNA-PRISM); goat-anti-rabbit-p1 and goat-anti-mouse-p12 (DNA-PRISM). Cells were incubated at room temperature for 1 h in the secondary antibody solution. Cells were washed 3 times with PBS, post-fixed for 15 min with 4% PFA and 4% wt/vol sucrose in PBS. This step was used to prevent cross-binding of the secondary antibodies to the primary antibodies in the following round of staining. Cells were washed 3 times with PBS and incubated again in the nuclear blocking buffer for 30 min at room temperature. Cells were then incubated overnight at 4 °C in the following primary antibody/peptide solution diluted in the nuclear blocking buffer for LNA- or DNA-PRISM: phalloidin-p2, Tuj-1-p3, cortactin-p4, SHANK3-p6, ARPC2-p7, bassoon-p8, synapsin-I-p9, Homer-1b/c-p10 (LNA-PRISM); phalloidin-p2, Tuj-1-p3, cortactin-p4, SHANK3-p6, bassoon-p5 (PNA), synapsin-I-p9, Homer-1b/c-p10 (DNA-PRISM). Cells were then washed 3 times with PBS. For LNA-PRISM, cells were incubated in diluted DAPI or Hoechst for 15 min. For DNA-PRISM, cells were incubated at room temperature for 1 h in donkey-anti-goat-Alexa488 diluted in the regular blocking buffer, washed 3 times with PBS, and then incubated with 10 nM of 100 nm diameter gold nanoparticle (Sigma-Aldrich) in PBS for 15 min. Cells were then washed 3 times with PBS prior to imaging. See **Table S1** for antibody information.

### Multiplexed confocal imaging of neurons using LNA-PRISM

LNA-PRISM imaging was performed on an Opera Phenix High-Content Screening System (PerkinElmer, Waltham, MA) equipped with a fast laser-based autofocus system, high NA water immersion objective (63×, numerical aperture=1.15), two large format scientific complementary metal-oxide semiconductor (sCMOS) cameras and spinning disk optics. 405 nm, 488 nm and 561 nm lasers were used as excitation for DAPI, MAP2, and VGLUT1 channels respectively. PRISM images were acquired using a 640nm laser (40 mW), sCMOS camera with 1–2 s exposure time, and effective pixel size of 187 nm. Before each imaging round, the corresponding imaging probe was freshly diluted to 10 nM in imaging buffer (500 mM NaCl in 1× PBS (Sigma-Aldrich), pH 8). Neurons were incubated with 10 nM imaging probes for 5 min, and then washed twice manually with imaging buffer to remove the free imaging probe. For each field, a stack of 3 images was acquired with a step of 0.5 μm. After imaging, cells were washed three times with wash buffer (0.01× PBS), and incubated in the wash buffer for 5 min after the last wash before the next round of imaging.

### LNA-PRISM confocal image processing and analysis

Lateral (x,y) drift between LNA-PRISM images from different imaging rounds was corrected by aligning the MAP2 channel in each imaging round. The (x,y) drift was estimated by locating the peak of the spatial cross-correlation function between two MAP2 images. Each image was first filtered using a top-hat filter with a disk structural element of 100 pixels to remove the uneven background in the image. For segmentation of synaptic punctae, the contrast of the image was first adjusted by saturating the highest and lowest 1% of pixels in the intensity histogram. The image was then denoised using a 5×5 Wiener filter, and filtered again with a top-hat operator with a disk structural element of 8 pixels to enhance the punctae in the image. The optimal threshold for each image was determined using an object-feature-based thresholding algorithm adapted from the thresholding algorithm previously used for single molecule tracking^42^. The threshold producing the maximum number of objects was chosen as the optimal threshold. We found thresholding based on the features of objects was more robust to the intensity variations across different channels than the intensity-based approach for synapse segmentation. Connected synapses in the thresholded image were then separated using a watershed transform to obtain the final segmentation mask for each image of each synaptic target. Synapses were identified using synapsin-I as the synapse marker following Micheva et al.^12^. Each segmented synapsin-I punctum larger than 0.42 μm^2^ was considered to be a synapse. For other synaptic proteins, only punctae that were colocalized (intensity weighted centroid distance < 1 μm) with synapsin-I punctae were considered to be synapses and therefore retained. For each identified synapse, the average intensity and area of the segmented punctum in each synaptic channel were measured; zero was assigned when no colocalized punctum was detected. For non-synaptic targets (MAP2 and Tuj-1), the intensity was estimated by averaging the intensity within the synapsin-I puncta and no area measurement was performed. The Pearson Correlation Coefficient between each pair of synaptic intensity measurement was computed for each cell culture batch. The average correlation coefficients over 3 cell culture batches (total 175,399 synapses) were represented using a network diagram. An edge was shown between two nodes if the corresponding correlation was greater than 0.39, with the thickness of each edge representing the strength of the correlation. t-Distributed Stochastic Neighbor Embedding (t-SNE) was used to visualize the distributions of synapses in high-dimensional feature space. Twenty features (intensity levels and punctae sizes of 10 proteins) of single synaptic profiles from a single replicate were used as the input to t-SNE. Each feature was normalized to have standard deviation of one and minimum of zero before applying t-SNE. t-SNE analysis was performed using scikit-learn 0.18.1 in Python 3.5 with the Barnes-Hut approximation, perplexity parameter equal to 100, and PCA initialization. The resulting t-SNE maps were similar for perplexity of 10-1000. Hierarchical clustering of single-synapse profiles was performed by first normalizing the distribution of each feature to have a minimum of zero and a standard deviation of one using all synapses as input. 24 features (intensity levels and punctae sizes of the 12 proteins) were used as input to the clustering analysis. Clustering was performed using the “clustergram” function in MATLAB R2015a (The MathWorks, Inc., Natick, MA) with the Euclidean metric and Ward’s linkage. The optimal number of clusters was determined using silhouette analysis.

### Multiplexed super-resolution imaging of neurons using DNA-PRISM

Single and dual channel PAINT imaging was performed on an inverted Nikon Eclipse Ti microscope (Nikon Instruments, Inc., Melville, NY) with the Perfect Focus System and oil-immersion objective (Plan Apo TIRF 100×, numerical aperture (NA) 1.49). A 642 nm wavelength laser (100 mW nominal) was used for excitation. Images were acquired using an Electron-Multiplying Charge-Coupled Device (EMCCD) camera (iXon DU-897, Andor Technology, Belfast, UK) with 100 ms exposure time, 100 EM gain, and effective pixel size of 160 nm. Nine-channel DNA-PAINT imaging was performed on an inverted Nikon Eclipse Ti microscope (Nikon Instruments) with the Perfect Focus System and oil-immersion objective (Plan Apo TIRF 100×, numerical aperture (NA) 1.49). 640 nm laser (45 mW nominal) was used for excitation. Images were acquired using a Zyla 4.2 sCMOS camera (Andor Technology) with 100 ms exposure time, 2×2 binning, and effective pixel size of 130 nm. Cells were imaged using Highly Inclined and Laminated Optical illumination (HILO). The same imaging probe sequences labeled with Atto655 (Eurofins, Luxembourg) and imaging/washing buffer (1× PBS (Sigma-Aldrich), 500 mM NaCl, pH 8) as previously published^23^ were used (**Table S2**). Probe exchange was performed using a home-built fluid control system (**Figure S14**). Depending on the labeling density, typically 0.5-3 nM imaging probe diluted in the imaging buffer was used in order to achieve optimal spot density for single molecule imaging. 5,000–20,000 image frames were typically acquired for each target.

### Super-resolution image reconstruction and localization analysis

Localization of the center of each diffraction-limited spot corresponding to a single molecule in the acquired movies was performed using DAOSTORM^43,44^. The super-resolution image was reconstructed by computing a 2D histogram of the (x,y) coordinates of the localized spots with bin size 5.4×5.4 nm, followed by smoothing with a 2D Gaussian filter. Non-specifically bound gold nanoparticles were used as fiduciary markers to estimate the drift and align images of distinct targets. Drift was estimated by fitting splines to the x- and y-coordinates separately of each fiduciary marker as a function of time using LOESS regression, and averaging over splines of all the fiduciary markers. 1D cross-sectional profiles of protein distributions were generated by projecting the 2D (x,y) coordinates of the localized spots onto the 1D coordinates along the trans-synaptic axes, and followed by computing a 1D histogram of the 1D coordinates. All image reconstruction and analysis procedures except for single molecule localization were performed using MATLAB R2015a (The MathWorks, Inc.).

## ACKNOWLEDGMENTS

Funding from NIH BRAIN Initiative Award 1U01MH10601101 to M.B. and E.S.B. is gratefully acknowledged. Funding from the NSF Physics of Living Systems program NSF PoLS 1305537 to M.B. is additionally acknowledged. E.S.B additionally acknowledges the HHMI-Simons Faculty Scholars Program, the Open Philanthropy Project, U. S. Army Research Laboratory and the U. S. Army Research Office under contract/grant number W911NF1510548, the New York Stem Cell Foundation-Robertson Award, and NIH grants 1RM1HG008525, 1R24MH106075, 1U01MH106011, and 1R01NS087950. J.R.C. acknowledges funding from the Stanley Center for Psychiatric Research. Dr. Ralf Jungmann is acknowledged for helpful discussions on DNA-PAINT at the outset of this work. We acknowledge the Koch Institute Swanson Biotechnology Center, specifically Microscopy and Biopolymers & Proteomics core facilities, as well as the Whitehead Institute W.M. Keck Microscopy Facility, for technical support.

## AUTHOR CONTRIBUTIONS

S-M.G., S.G., R.V., E.S.B., and M.B conceived of and designed the DNA-PRISM study. S-M.G. and M.B. conceived of the LNA-PRISM study. S-M.G., L.L., J.R.C., and M.B. designed the LNA-PRISM study. R.V. designed and implemented the antibody conjugation protocols for DNA- and LNA-PRISM. S-M.G and S.G. designed and implemented the DNA-PRISM experiments. S-M.G analyzed the DNA-PRISM data. D.P. and L.L. prepared primary neuronal cultures. S-M.G., S.G., and R.V. designed and implemented the DNA-PRISM fixation, blocking, staining, probe exchange, and imaging protocols. S-M.G. and L.L. adapted the DNA-PRISM protocol for LNA-PRISM. S-M.G., L.L., and S.G. analyzed the LNA-PRISM data. S.G., A.B.K, and P.C.B. designed the microfluidic device for DNA-PRISM. M.B. designed and supervised the overall project including DNA-PRISM and LNA-PRISM. J.R.C. supervised the LNA-PRISM project. S-M.G., S.G., and M.B. interpreted the DNA-PRISM results. S-M.G., S.G., L.L., J.R.C., and M.B. interpreted the LNA-PRISM results. S-M.G, S.G., and M.B. wrote the manuscript. All authors commented on the manuscript.

